# Membrane tubulation by adhesion of spherical nanoparticles

**DOI:** 10.64898/2026.02.17.706332

**Authors:** Thomas R. Weikl

## Abstract

Adhesion of spherical nanoparticles or virus-like particles to membranes can lead to membrane tubules in which linear chains of adhering particles are cooperatively wrapped by the membrane. This cooperative wrapping of spherical particles in tubules is energetically favourable compared to the individual wrapping of the particles because of a favourable interplay of bending and adhesion energies in the contact regions in which the membrane detaches from the particles, and because a particle in a tubule has two such contact regions in the membrane necks that connect the particle to the neighbouring particles, whereas an individually wrapped particle has only one contact region to the surrounding membrane. The energetic gain of cooperative wrapping strongly depends on the range of the particle-membrane adhesion potential, which determines the size of the contact regions. At sufficiently large adhesion energies for wrapping, the energy gain Δ*E* per particle is only weakly affected by the membrane tension *τ* as long as the characteristic length 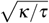 of the membranes with bending rigidity *κ* is clearly larger than the contact regions. For large particle adhesion energies at which the particles are fully wrapped, however, Δ*E* can be limited by the minimum possible radius of the membrane necks, depending on the adhesion potential range.

## 1 Introduction

Cellular uptake of nanoparticles can occur via a variety of endocytic pathways ^1,2^, which all require the generation of cell membrane curvature and membrane invaginations. Nanoparticles that adhere to cell membranes can themselves generate membrane curvature if their adhesion energy is sufficiently large to compensate for the cost of membrane bending ^3^. The nanoparticles then can be partially or fully wrapped by the membrane, depending on their shape and size ^4,5^, the bending rigidity and tension of the membrane ^6^, the energy and range of the adhesion potential ^7^, and the curvature of the plasma membrane segment or membrane vesicle prior to wrapping ^8,9^ in the interplay of the elastic energy of the membrane and the adhesion energy of the particles.

Nanoparticles can also be cooperatively wrapped as linear chains of particles in membrane tubules, besides being wrapped individually as single particles. In computational approaches, cooperative wrapping of spherical nanoparticles in membrane tubules has been both found in Monte Carlo simulations in which the membranes are modelled as triangulated elastic surfaces ^10,11^ and in coarse-grained molecular simulations of particle-membrane systems ^12–16^. In experiments, cooperative wrapping in membrane tubules has been observed for spherical citrate-stabilized gold nanoparticles with a diameter of 10 nm in contact with POPC or DOPC large unilamellar vesicles ^17^ (see Figure 1a). The gold nanoparticles are arranged linearly with a distance of about 3 nm in the membrane tubules, which protrude into the vesicle interior and exhibit periodic shaping of the membrane around the wrapped particles ^17^. Particle-induced membrane tubulation of cells has been first reported for spherical virus-like particles with a diameter of 45 nm that are composed of capsid proteins binding to GM1 glycolipids in the cell membranes (see Figure 1c) ^18^. The particles are linearly arranged in the cell membrane tubules, with varying distances between the particles (see arrows in Figure 1c). More recently, the role of the particle adhesion energy for cell membrane tubulation has been explored in an artificial system of capsid particles that are densely covered with green fluorescent proteins (GFPs) and bind to anti-GFP nanobodies anchored to the cell membranes (see Figure 1d) ^19^. In this system, the particle adhesion energy was varied by using a repertoire of anti-GFP nanobodies with different affinities for GFP. For nanobodies with affinities beyond a threshold value, membrane tubulation was observed both for energy-depleted cells and giant unilamellar vesicles (GUVs) in fluorescence micrographs. High-resolution electron microscopy images correlated with fluorescence recordings indicate membrane tubules filled by particle chains (see Figure 1e). Pairs of rod-shaped particles with a length of several micrometers have been observed to be cooperatively wrapped in tubular membrane invaginations of GUVs with a tip-to-tip orientation of the particles ^20^. In this system, particle adhesion to GUVs was induced by depletion interactions from nonadsorbing polyacrylamide polymers in the solvent. For micron-sized spherical particles adhering to GUVs via specific biotin-avidin repectorligand interactions, tube-like structures in which two particles are cooperatively wrapped have been observed as a rather rare conformation among other dimer conformations ^21,22^.

**Fig. 1.**
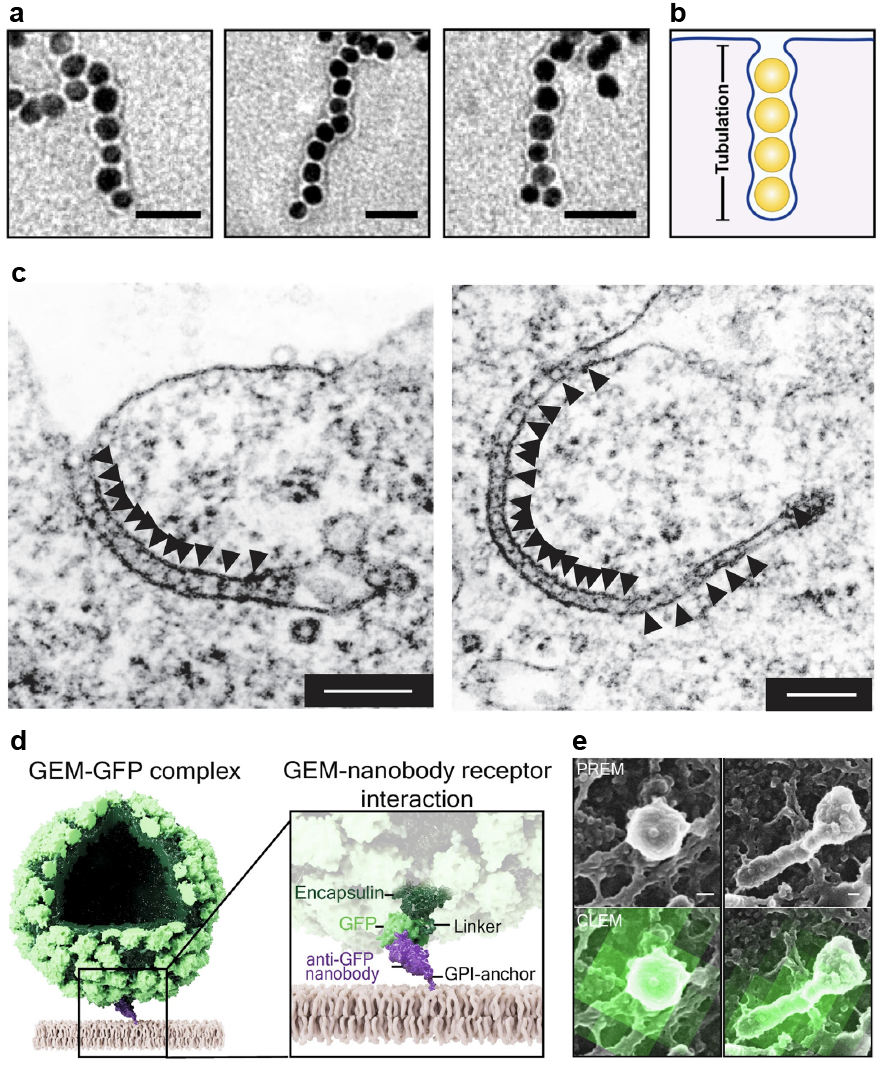
(a) Linear chains of citrate-stabilized gold nanoparticles with a diameter of about 10 nm in membrane tubules of large unilamellar vesicles. The scale bars of the transmission electron microscopy pictures are 50 nm. The particle-filled membrane tubules protrude into the vesicle interior as sketched in (b). Reproduced with permission from Ref. 17. (c) Linear chains of virus-like particles with a diameter of 45 nm (highlighted by triangular arrows) in tubular membrane invaginations of cells. The scale bar of the electron micrographs are 200 nm. The particles are composed of 72 icosahedrally organized capsid protein pentamers that contain a binding site for cell membrane GM1 glycolipids in each monomer. Control experiments indicate that the membrane invaginations are induced by particle adhesion without the help of active endocytic machinery or caveolar coats ^18^. Reproduced with permission from Ref. 18. (d) Illustration of a genetically encoded nanoparticle (GEM) assembled from encapsulin proteins in the artificial adhesion system of Ref. 19. The particles are densely covered by GFP proteins coupled to the encapsulin proteins and bind to anti-GFP nanobodies anchored to the cell membranes. (e) Platinum-replica electron microscopy (PREM) and correlative light electron microscopy (CLEM) pictures of plasma membrane sheets generated after unroofing cells in contact with GEMs. The elongated green fluorescent structure on the right indicates a short GEM-filled membrane nanotube. The GEMs are not wrapped and internalized by endocytic events involving chlathrin-coated pits or caveolae according to control experiments. The scale bars are 50 nm. Reproduced with permission from Ref. 19.

The cooperative wapping of nanoparticles observed in these computational and experimental systems implies an energy gain for the wrapping of linear chains of particles in membrane tubules, compared to the individual wrapping of the particles as single particles. For spherical nanoparticles, the energy gain of cooperative wrapping in membrane tubules has been found to depend strongly on the interaction range of the adhesion potential relative to the particle radius, and to vanish if the interaction range becomes negligibly small ^7^. In energetic arguments ^3,8^, in energy calculations based on the shape equations of rotationally symmetric membrane vesicles ^6,9^, and in energy minimizations ^23^ with the popular “Surface Evolver” ^24^ software, the interaction range of particles adhering to membranes is often assumed to be zero out of computational convenience or necessity, because a zero potential range allows a sharp distinction between adhering and non-adhering membrane segments. However, the assumption of zero potential range is only a valid approximation in determining the membrane-wrapping behavior of spherical particles if the particle radius is at least three orders of magnitude larger than the interaction range of the particle-membrane adhesion potential ^7^, so, e.g., for micron-sized particles with short-ranged adhesive interactions on the length scale of a nanometer. For a potential range of zero, spherical particles are fully wrapped by a tensionless, initially planar membranes as soon as the bending energy cost 8*πκ* of full wrapping is compensated by the adhesion energy 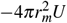 where *κ* is the bending rigidity of the membrane, −*U* is the adhesion energy per area, and *r*_*m*_ is the radius of the spherical membrane vesicle wrapping the particle ^3,25^. The spherical vesicle around the particle is connected to the surrounding planar membrane by a membrane neck of catenoidal shape, which does not contribute to the overall bending energy because the mean curvature of the catenoid is zero. Similarly, energy calculations for wrapping chains of particles at zero range of the particle-membrane adhesion potential predict full wrapping of the particles beyond the threshold adhesion energy 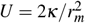, with catenoidal necks connecting the particles ^25^. At zero potential range, the overall energy 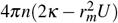 of wrapping *n* particles individually by a tensionless membrane or cooperatively as chain of particles therefore is the same. This calculation implies that also the membrane thickness and, thus, the catenoidal neck limited by this thickness, is negligibly small compared to the vesicle membrane radius *r*_*m*_.

At realistic finite potential ranges, in contrast, adhering and non-adhering membrane segments of a spherical nanoparticle are separated by a finite contact region in which the membrane detaches from the particle. In this contact region, the membrane already “swings into” the catenoidal shape of the surrounding non-adhering membrane, but still gains adhesion energy, which leads to a favourable contribution to the overall energy of wrapping ^7^. The energy gain for the cooperative wrapping of spherical particles in membrane tubules then simply results from the fact that a central particle in a membrane tubule has two of these energetically favourable contact regions in the membrane necks to the neighbouring particles, compared to a single contact region for an individually wrapped particle in the neck to the surrounding non-adhering membrane (see Figure 2).

**Fig. 2.**
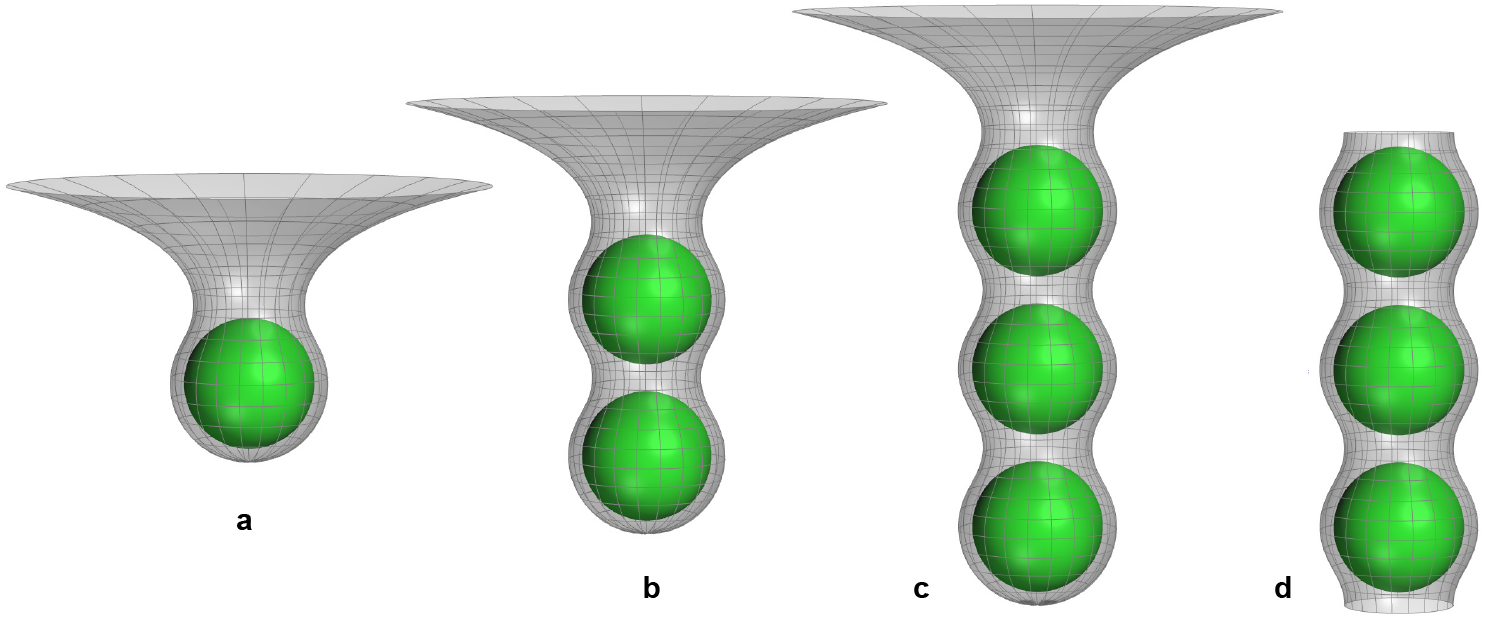
Minimum-energy conformations of (a) a single spherical particle, (b) two particles, (c) three particles, and (d) three central particles in a linear chain of many particles wrapped by a tensionless membrane for the rescaled adhesion energy *u* = 2.5, particle radius *r*_*p*_ = 19 nm, membrane radius *r*_*m*_ = 23 nm, and standard deviation *σ* = 1 nm of the particle membrane adhesion potential (2). The rotationally symmetric membrane shapes in (b) and (c) have been constructed from segments of the numerically determined shapes in (a) and (d).

For elongated particles, e.g. prolate or sphero-cylindrical, rod-shaped particles, cooperative wrapping in membrane tubules with tip-to-tip orientation provides an energy gain compared to the individual wrapping of the particles also for negligibly small ranges of the adhesion potential ^26^. For these particles, the wrapping of the highly curved tips costs more bending energy per area than the wrapping of the less curved sides, and the wrapping in membrane tubules is energetically favourable because both tips of a central particle in the tubule remain unwrapped. In line with these predictions based on minimizations of the sum of bending and adhesion energies, micron-sized rod-shaped particles have been observed to have a strong tendency to be cooperatively wrapped tip-to-tip by membrane tubules ^20^, whereas micron-sized spherical particles with an adhesion potential range that is several orders of magnitude smaller than the particle diameter exhibit tube-like structures in which two particles are cooperatively wrapped only as a rather rare conformation among other dimer conformations ^21,22^. For spherocylindrical nanoparticles, cooperative wrapping by membrane tubules with tip-to-tip orientation of the particles as also been reproduced in coarse-grained molecular simulations ^27^.

In this article, we revisit and extend the elastic model of Ref. 7 for the cooperative wrapping of spherical particles in tubules (i) by including membrane tension and (ii) by considering the minimum radius of membrane necks and the particle radius *r*_*p*_ as length parameters in addition to the adhesion potential range and the membrane vesicle radius *r*_*m*_. For a fully wrapped spherical particle, the membrane tension *τ* leads to the energy contribution 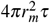 that needs to be compensated by the adhesion energy of the particle, in addition to the bending energy cost 8*πκ* of full wrapping (see above). The membrane tension *τ* therefore increases the adhesion energy required for particle wrapping ^6^. At sufficiently large adhesion energies for wrapping, we find that the energy gain Δ*E* of cooperative wrapping per particle is only weakly affected by the membrane tension *τ* as long as the characteristic length *κ/τ* of the membranes with bending rigidity *κ* is clearly larger than the contact regions in which the membrane detaches from the particles. For large particle adhesion energies at which the particles are fully wrapped, the energy gain Δ*E* of cooperative wrapping can be limited by the minimum possible radius of membrane necks, depending on the adhesion potential range. The particle radius *r*_*p*_, in contrast, affects the conformation and energy of particle-filled tubules only at small adhesion energies at which the particles are in direct contact in the tubules with distance *d* = 2*r*_*p*_.

## 2 Methods

### 2.1 Model

The membrane curvature generation and wrapping of adhesive particles results from the interplay of the elastic energy of the membrane and the adhesion energy of the particles. The total energy *E* is the sum

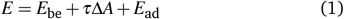

of the bending energy *E*_be_ = 2*κ M*^2^ dA with local mean curvature *M*, the adhesion energy *E*_ad_ of the particles, and the energy *τ*Δ*A* associated with the membrane tension *τ*, where Δ*A* is the excess area of the curved membrane relative to initially planar membrane prior to particle adhesion.

We describe the particle-membrane adhesion potential by the Gaussian potential

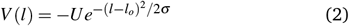

where −*U* is the adhesion energy per area, *l*_*o*_ is the preferred binding separation of the membrane midplane from the particle surface, and *ξ* is the standard deviation of the Gaussian function that reflects the range of the adhesion potential. The Gaussian potential (2) is plausible if the particle adhesion is mediated by receptor and ligand molecules that are flexible anchored to the particles and the membrane ^28^. The preferred binding separation *l*_*o*_ and standard deviation *ξ* of the adhesion potential then depend on the dimensions of the receptor-ligand complex, which can tilt due to the flexible anchoring, and the lengths of the flexible linkers that anchor the receptors and ligands, and the adhesion energy −*U* per area depends on the ligand density on the particle surface and the binding constant of the receptor-ligand interaction ^19^ (see Eq. (5) below).

Three central length parameters of the model are the particle radius *r*_*p*_, the radius *r*_*m*_ = *r*_*p*_ + *l*_*o*_ of membrane segments that wrap the particle at the preferred binding separation *l*_*o*_, and the standard deviation *σ* of the adhesion potential (2). Besides the membrane radius *r*_*m*_, the particle radius *r*_*p*_ is a relevant parameter for the cooperative wrapping in membrane tubules because it limits the distance *d* ≥ 2*r*_*p*_ of particles in tubules. A fourth length scale of the model is the minimum membrane-membrane distance in membrane necks.

Besides these length parameters, two central dimensionless parameters of the model are the rescaled adhesion energy

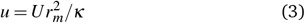

and the rescaled tension

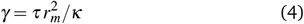

which can be derived from the three energetic parameters *U, κ*, and *τ* by understanding the bending rigidity *κ* as energy unit and the membrane radius *r*_*m*_ as length unit in approaches focussing on minimum total energies as here.

To limit the number of independent parameters, we initially focus on length parameters adapted from a particle system previously presented and modelled in Ref. 19 (see Fig. 1d). In this system, a capsid particle with a radius of 15 nm is densely covered by 180 green fluorescent proteins (GFPs), which are connected by unstructured 12-residue peptide linkers to the capsid proteins, leading to a radius *r*_*p*_ = 19 nm of the GFP-covered particle. The GFPs bind to anti-GFP nanobodies attached to the cell membranes by glycosylphosphatidylinositol (GPI) anchors. Based on the dimensions and linker attachment sites of the GFP-nanobody complex and the flexibility of the linkers, which allow also for tilting of the complex, the adhesion potential was estimated as the Gaussian potential (2) with standard deviation *σ* = 1 nm and preferred binding separation *l*_*o*_ = 8 nm between capsid particle surface and membrane midplane ^19^. A membrane vesicle wrapping a single particle thus has the radius *r*_*m*_ = (15 + 8) nm = 23 nm. In this continuum modeling of particle adhesion, the adhesion energy per area ^19^

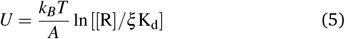

depends on the vesicle area per protein complex 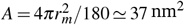, the area concentration [R] of GPI-anchored nanobodies in the cell membrane, and the dissociation constant *K*_*d*_ of the GFP-nanobody complex. The parameter *ξ* in this equation is a “conversion length” from 3D binding of the soluble complex to the 2D binding of the anchored complex at the preferred separation *l*_*o*_^29^.

### 2.2 Minimization method

To determine the minimum-energy shapes of the rotationally symmetric membranes around individually wrapped particles or co-operatively wrapped linear chains of particles, we extend the methodology of Ref. 7 to membranes under tension. In this methodology, we discretize the profiles of the rotationally symmetric membrane shapes in two different parametrizations.

In *parametrization 1*, the rotationally symmetric membrane shapes are described by the function *r*(*z*) where *z* is the coordinate along the axis of rotation, and *r* is the radial distance from this axis. In this parametrization, the bending energy *E*_be_, membrane area *A*, and adhesion energy *E*_ad_ can be expressed as

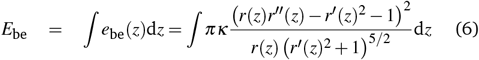

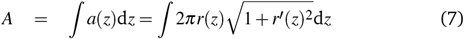

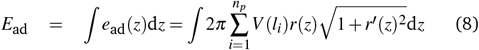

with *l*_*i*_ = *l*_*i*_(*z, r*(*z*)) and the bending energy density *e*_be_(*z*), area density *a*(*z*), and adhesion energy density *e*_ad_(*z*). Here, primes indicate derivatives with respect to *z*, and *n*_*p*_ is the number of considered particles, which are centered along the *z*-axis. For a membrane segment adhering at preferred separation *l*_*o*_ to a particle centered at *z* = 0, the membrane profile is 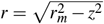, and the densities are *e*_be_(*z*) = 4*πκ/r*_*m*_, *a*(*z*) = 2*πr*_*m*_, and *e*_ad_(*z*) = −2*πr*_*m*_*U*, which are constant functions independent of *z*. If the particle is fully wrapped by the membrane from *z* = −*r*_*m*_ to *z* = *r*_*m*_, these densities lead to 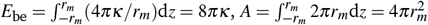, and 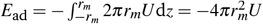 as expected (see above).

We use parametrization 1 to determine the membrane shapes around central particles in membrane tubules with distance *d* ≥ 2*r*_*p*_ between the particles. Because of the periodicity of the tubular shapes (see Fig. 2), we determine the minimum energy shape of a membrane segment from the center of a particle to the center of the membrane neck that connects to the neighbouring particle in the tubule. We discretize the membrane profile *r*(*z*) of this membrane segment using 400 discretization points, express the derivates *r′*(*z*) and *r′′*(*z*) as standard finite differences, and minimize the total energy *E* with respect to the radial distances *r*(*z*_*i*_) at the discretization points *z*_*i*_ and with respect to the distance *d* of particles using the software Mathematica 14.3^30^. The number of particles in this minimization is *n*_*p*_ = 2, because only the two particles connected by the membrane neck of the segment contribute to the adhesion energy of the segment. For central particles in the tubule, the excess area Δ*A* of the membrane segments wrapping these particles relative to the planar membrane prior to tubule formation is equal to the area *A* of the segments.

We also use parametrization 1 to determine the rotationally symmetric membrane shape around a single, deeply wrapped particle, e.g. the left shape of Figure 2. A complication here is that the derivative *r′*(*z*) diverges at the membrane point on the symmetry axis, i.e. the point at the bottom of the left shape of Figure 2, and when the non-adhering membrane approaches the surrounding planar membrane. The divergence of *r′*(*z*) at the membrane point on the symmetry axis can be treated by taking into account that the energy densities *e*_be_(*z*), *e*_ad_(*z*) and *τa*(*z*) are constant functions for the adhering membrane segment with spherical membrane shape (see above). The divergence of *r′*(*z*) as the non-adhering membrane approaches the surrounding planar membrane, in contrast, is unproblematic only in the case of zero membrane tension, because the non-adhering membrane then adopts a catenoidal shape with zero total energy because the bending energy of the catenoid is zero. For tensionless membranes, the membrane profile therefore just needs to be determined up to a discretization point *z*_*n*_ of the non-adhering membrane at which the total energy is zero ^7^. For membranes with finite tension, we numerically determine the membrane profile also up to a discretization point *z*_*n*_ of the non-adhering membrane around a single deeply wrapped particle at which the slope *r′*(*z*_*n*_) still allows a reliable numerical determination of the total energy. For *z* values beyond this point, we use the analytical solution 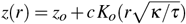 with Bessel function *K*_*o*_ for rotationally symmetric membrane shapes with small gradients *z′*(*r*) ^31^, which we fit to the last 20 points of the numerically determined profile until point *z*_*n*_. In the energetic data presented in the Results section, the analytical continuation of the non-adhering membrane shape contributes only marginally to the total energy. In the numerical minimization of the total energy of the discretized membrane profile, we use a discretization length *δz* = *r*_*m*_*/*400 and up to a total of 1000 discretization points, depending on the location of the point *z*_*n*_. The excess area of the discretized membrane up to *z*_*n*_ is determined as Δ*A* = *A*− 2*πr*(*z*_*n*_)^2^ with area *A* calculated based on Eq. (7).

To determine to rotationally symmetric membrane shape around particles that are less than half-wrapped by the membrane, we use parametrization 2 in which the membrane profile is described by the function *z*(*r*). In this parameterization, the bending energy *E*_be_, excess area Δ*A*, and adhesion energy *E*_ad_ can be expressed as

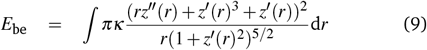

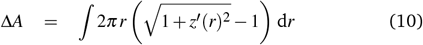

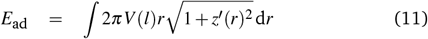

with *l* = *l*(*z*(*r*), *r*). To numerically determine the membrane profile *z*(*r*) by minimisation of the total energy, we use the discretization length *δr* = *r*_*m*_*/*400 and up to 3000 discretization points in numerical minimizations with Mathematica 14.3.

## 3 Results

We first consider the cooperative wrapping of linear chains of particles in membrane tubules, with a focus on how the rotationally symmetric minimum-energy membranes shapes and particle distances of the tubules depend on the rescaled membrane tension *γ* and the rescaled adhesion energy *u* of the particles defined in eqs. (3) and (4). We initially adjust the length parameters of our model to the particle system of Fig. 1d (see Methods), but consider later the effect of these length parameter values on the results. Figure 3 illustrates how the conformations of the particle-filled membrane tubules depend on the rescaled adhesion energy *u* for tensionless membranes with *γ* = 0 already considered in Ref. 19 and for membranes with rescaled tension *γ* = 0.2, 0.5, and 1. At the value 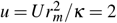 at which the adhesion energy 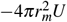 of a spherical vesicle with radius *r*_*m*_ fully wrapping a single particle at preferred separation *l*_*o*_ of the adhesion potential (2) is oppositely equal to the bending energy 8*πκ* of the vesicle, the particles are in contact in the tubule with distance *d* = 2*r*_*p*_ between the particles (see Fig. 3c), and the membrane tubules are only weakly undulated with a neck radius *r*_*n*_ larger than 0.8*r*_*m*_. Here, *r*_*m*_ is the membrane-midplane radius of membrane segments wrapping the particle at preferred separation *l*_*o*_ and, thus, the maximum radius of the tubules at cross sections that contain the particle centres. For *γ* = 0 and 0.2, the tubule conformations change continuously with increasing *u* (see Fig. 3a for *γ* = 0.2). At these values of *γ*, the particle distance *d* and tubule neck radius *r*_*n*_ start to increase continuously with *u* at the transition values *u*_*t*_ ≃ 2.41 for *γ* = 0 and *u*_*t*_ ≃ 2.69 for *γ* = 0.2. At large values of *u*, the tubule conformations are strongly undulated and attain the minimum neck radius *r*_*n*_ = 2.5 nm for the membrane thickness 5 nm assumed here (gray dashed line at *r*_*n*_*/r*_*m*_ = 2.5*/*23 ≃ 0.109 in Fig. 2d). At the minimum neck radius, the neck is closed, i.e. the membrane segments lining the neck are in contact. At large values of *u*, the particle distance tends to *d* ≃ 2.75*r*_*p*_, with a distance maximum at intermediate values around *u* = 3.3 (see Fig. 3d). For *γ* = 0.5 and 1, in contrast, the tubule conformations change discontinuously at the transition values *u*_*t*_ ≃ 3.09 and 3.54, respectively. At these transition values, the tubule conformation changes abruptly from weakly undulated to deeply undulated as illustrated in Fig. 3b for *γ* = 1.

**Fig. 3.**
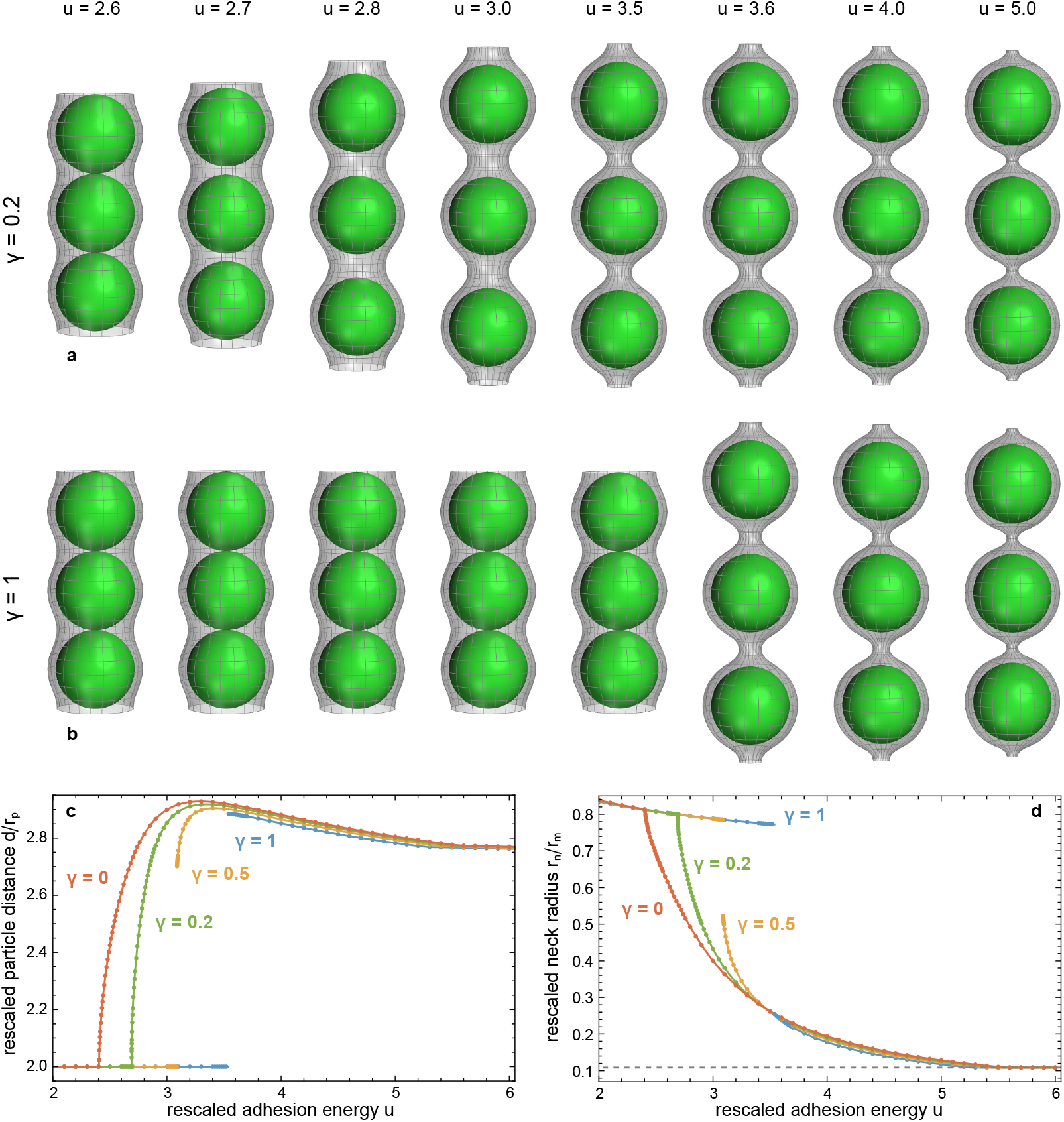
(a) and (b) Minimum-energy conformations of three central particles in a membrane tubule wrapping many particles at the rescaled membrane tension *γ* = 0.2 and 1 at various values of the rescaled adhesion energy *u*. (c) Particle distance *d* and (d) neck radius *r*_*n*_ in a membrane tubule wrapping many particles versus rescaled adhesion energy *u* at the rescaled membrane tensions *γ* = 0, 0.2, 0.5, and 1. The particle radius here is taken to be *r*_*p*_ = 19 nm, the preferred distance of adhering membrane segments from the particle center is *r*_*m*_ = 23 nm, and the standard deviation of the particle membrane adhesion potential (2) is *σ* = 1 nm (see Methods). The minimum neck radius of 2.5 nm for a membrane thickness of 5 nm is represented by the gray dashed line in (d).

Cooperative wrapping of linear chains of particles in membrane tubules requires an energetic gain compared to the individual wrapping of the particles. In Fig. 4a, the minimum total energy *E* per central particle of a long tubule is compared to the minimum total energy *E* of an individually wrapped particle. The data points in Fig. 4 represent minimization results, and the lines are interpolations of the data points. For particles in tubules, and for individually wrapped particles in tensionless membranes with *γ* = 0, the results obtained with our minimization methods are numerically precise at all values of the rescaled adhesion energy *u*. For individually wrapped particles in membranes under tension, the minimization methods are (i) numerically precise for particles that are less than half wrapped and (ii) numerically reliable for deeply wrapped particles with large adhesion energy *u*. At intermediate values of *u* at which no minimization results are available, we interpolate the energies from the results obtained in the regimes (i) and (ii). These interpolations benefit from the linear dependence of the energy *E* on large values of *u* in regime (ii). In regime (i), we fit the minimization data points by a cubic function, which is extended in the dashed interpolation lines in Fig. 4 for *γ >* 0 until the intersection point with the linear fit obtained in regime (ii).

**Fig. 4.**
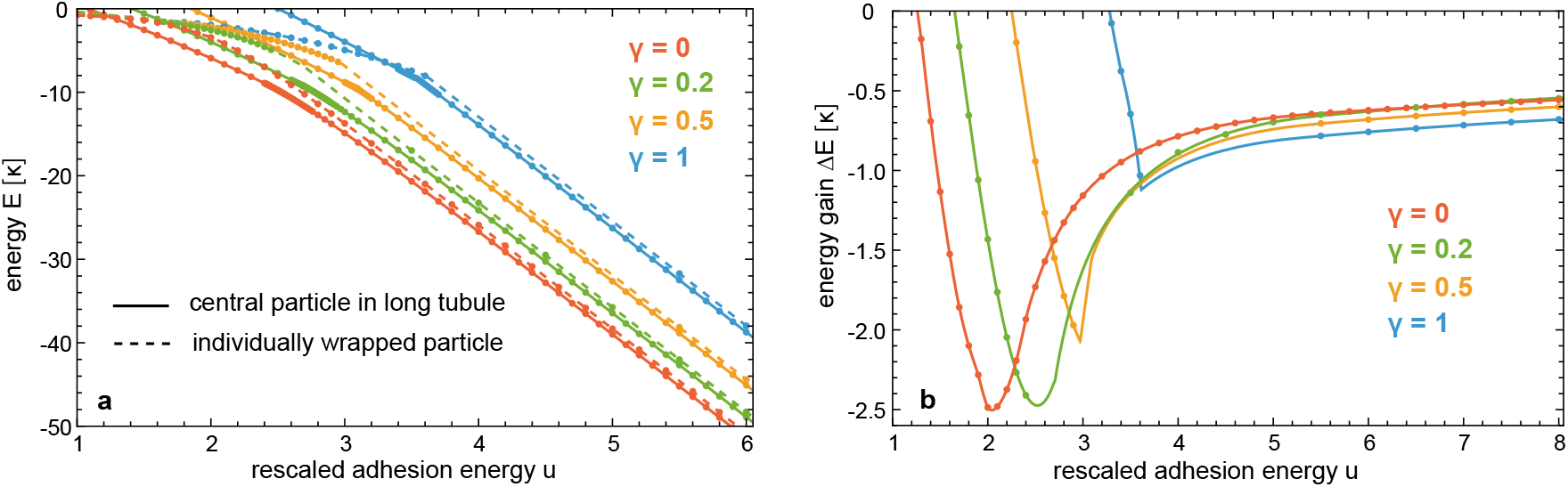
(a) Total energy *E* of individually wrapped particles (points interpolated by dashed lines) and per central particle in long membrane tubule wrapping many particles (points interpolated by full lines) versus rescaled adhesion *u* at different values of the rescaled tension *γ* for the same parameter values as in Figure 3. (b) Energy gain Δ*E* per central particle in long membrane tubule relative to an individually wrapped particle versus rescaled adhesion *u* at different values of *γ*.

At sufficiently large rescaled adhesion energies *u*, the minimum total energy *E* per central particle of a long tubule is lower than the minimum total energy *E* of an individually wrapped particle (see Figure 4a). The resulting energy gain Δ*E* of cooperative wrapping, defined as difference of the two energies of Fig. 4a, is illustrated in Fig. 4b. For tensionless membranes with *γ* = 0, the energy gain Δ*E* of cooperative wrapping, relative to individual wrapping, is maximal close to *u* = 2 at which the adhesion energy of membrane segments adhering at preferred separation *l*_*o*_ of the adhesion potential (2) is oppositely equal to the bending energy of these segments (see above). At the value *u* = 2 of the rescaled adhesion energy, single particles are half wrapped by a tensionless membrane ^7^. With increasing rescaled membrane tension *γ*, the energy Δ*E* attains its maximal value are larger and larger values of *u*, because the membrane tension impedes wrapping. At large values of *u*, the energy gain Δ*E* of membranes under tension is rather close to the energy Δ*E* for a tensionless membrane, in particular at the rescaled tensions *γ* = 0.2 and 0.5. At these large values of *u*, individual particles as well as particles in tubules are rather deeply wrapped by the membrane, and connected to the non-adhering membrane and the adjacent particles in the tubule, respectively, by membrane necks (see Figures 2 and 3). As previously shown for tensionless membranes ^7^, the energy gain for the cooperative wrapping in tubules results from a favourable interplay of bending and adhesion energies in the contact regions in which the membrane detaches from the particle. In this contact region, the membrane already “swings” into the catenoidal shape of the tensionless membrane neck, but still gains adhesion energy because of the finite range of the adhesion potential. The energy gain of cooperative wrapping then simply results from the fact that a central particle in a membrane tubule has two of these energetically favourable contact regions in the membrane necks to the two neighbouring particles, compared to a single neck and, thus, single contact region of an individually wrapped particle.

Figure 5b compares the total energy densities *e* of a central particle in a tubule and of an individually wrapped particle for the rescaled adhesion energy *u* = 5 at the values 0, 0.2, 0.5, and 1 of the rescaled tension *γ* considered here. The total energies *E* of Figure 4a result from an integration of these energy densities (see Methods). As functions of the rescaled coordinate *z/r*_*m*_, the total energy density of spherical membrane segments with radius *r*_*m*_ attains the constant values *e* = −2*πκ*(3 −*γ*) at *u* = 5. For individually wrapped particles centered at *z/r*_*m*_ = 0, the total energy densities in Figure 5c (dashed lines) adopt this value in the adhering membrane segment from *z/r*_*m*_ = −1 to about *z/r*_*m*_ = 0.5. For a particle located at *z/r*_*m*_ = 0 in a membrane tubule, the total energy densities in Figure 5c (full lines) are symmetric around *z/r*_*m*_ = 0 and adopt the value *e* = −2*πκ*(3 −*γ*) of a spherical membrane segment with radius *r*_*m*_ between about *z/r*_*m*_ = −0.5 and *z/r*_*m*_ = 0.5. For *z/r*_*m*_ *>* 0.5, the total energy densities of an individually wrapped particle and a central particle in a tubule are closely similar due to similar membrane shapes in the contact regions in which the membrane detaches from the particle and forms a neck (see Figure 5a and b for exemplary membrane shape profiles at *γ* = 0.5).

**Fig. 5.**
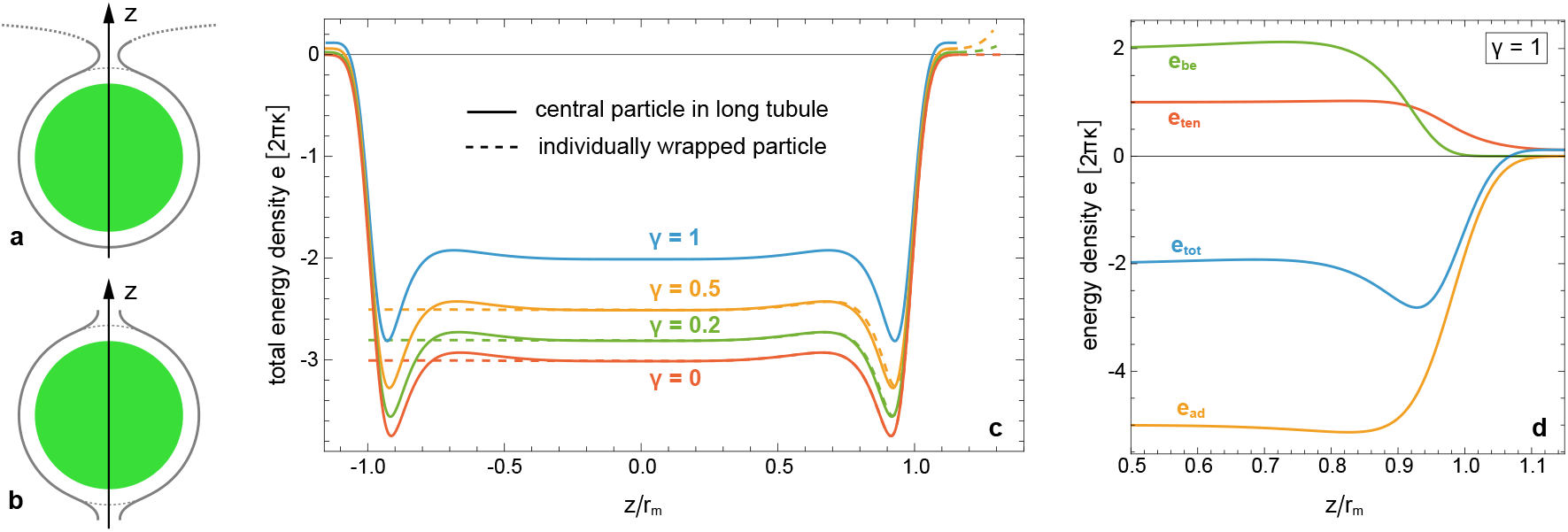
Membrane shape profiles and energy densities at the rescaled adhesion energy *u* = 5. (a) Membrane shape profile for an individually wrapped particle and (b) membrane shape profile for a central particle in a tubule at the rescaled tension *γ* = 0.5. The dotted continuation of the membrane neck in (a) results from a fit of the analytical solution 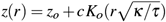 of the shape equation for small gradients *z*^*′*^(*r*) to the numerically determined shape (see Methods). The thin dashed lines in (a) and (b) represent a circle with membrane radius *r*_*m*_ = 23 nm. (c) Total energy densities *e* of a central particle in a tubule (full lines) and an individually wrapped particle (dashed lines) centered at *z* = 0 versus rescaled coordinate *z/r*_*m*_ at various values of the rescaled tension *γ*. (d) Bending energy density *e*_be_, adhesion energy density *e*_ad_, energy density *e*_ten_ associated with the membrane tension, and total energy density *e* = *e*_be_ + *e*_ten_ + *e*_ad_ of a central particle in a tubule for values of the rescaled coordinate *z/r*_*m*_ ≥ 0.5 at the rescaled tension *γ* = 1. In adhering membrane segments with radius *r*_*m*_, these energy densities attain the constant values *e*_be_ = 4*πκ, e*_ad_ = −2*π*^2^U = −2*πκ*u = −10*πκ* for *u* = 5, *e*_ten_ = 2*πrm*^2^*τ* = 2*πκγ* = 2*πκ* for *γ* = 1, and *e* = −4*πκ*. In the contact region in which the membrane detaches from the particle, the total energy density *e* attains a minimum at about *z/r*_*m*_ = 0.9, because the bending energy density *e*_be_ drops to zero at smaller values of *z/r*_*m*_ than the adhesion energy density *e*_ad_. In the contact region, the membrane shape changes from the spherical shape of the adhering membrane segment with radius *r*_*m*_ to the catenoidal shape of the membrane neck with bending energy zero, but still gains adhesion energy because of the finite range of the adhesion potential (2) with standard deviation *σ* = 1 nm. The shape profiles are not affected by the particle radius *r*_*p*_ and the minimum neck radius *r*_*n*_ = 2.5 nm because the particles are not in contact in the tubules and because the membrane necks are not yet closed at *u* = 5 (see Figure 3c and d).

The total energy densities first slightly increase for *z/r*_*m*_ *>* 0.5, then adopt a minimum around *z/r*_*m*_ = 0.9, and finally attain a value of 0 for a tensionless membrane with *γ* = 0 or positive values for *γ >* 0 in the non-adhering membrane neck that connects the wrapped particle to the surrounding planar membrane or the neighbouring particle in the tubule. The energy densities in Figure 5d show that the initial increase of the total energy density for *z/r*_*m*_ *>* 0.5 results from an increase in the bending energy density *e*_be_, which then drops to zero in the membrane neck, indicating that the neck attains a catenoidal shape even at the largest rescaled tension *γ* = 1 considered here. The adhesion energy density drops to zero at larger values of *z/r*_*m*_ than the bending energy density, which leads to the minimum of the total energy density in the contact region. The energy densities of Figure 5 are not affected by the minimum neck radius *r*_*n*_ = 2.5 nm because the membrane necks are not yet fully closed at the rescaled adhesion energy *u* = 5 for the standard deviation *σ* = 1 nm of the adhesion potential (2) considered here.

An energy gain Δ*E* of cooperative wrapping in tubules implies that the total energy contribution of the contact region is favourable. For *γ* = 0, the energy densities for a central particle in a tubule and an individually wrapped particle are identical in the region *z/r*_*m*_ *>* 0 that includes the contact region of the individually wrapped particle and one of the contact regions of the particle in the tubule. The energy gain Δ*E* thus results from the differences in the total energy densities for *z/r*_*m*_ *<* 0. In this region, the individual particle is wrapped by a spherical membrane segment with radius *r*_*m*_, with total energy contribution 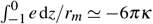. For the central particle in the tubule, the total energy density for *z/r*_*m*_ *<* 0 is symmetric to the profile for *z/r*_*m*_ *>* 0 and contributes 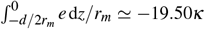, which is smaller than −6*πκ* and leads to the energy gain Δ*E* ≃ (−19.50 + 6*π*)*κ* ≃ −0.65*κ*. For *γ >* 0, the energy gain Δ*E* results from the differences of the total energies *E* obtained by integration of the whole energy densities, because the energy densities slightly differ also for *z/r*_*m*_ *>* 0.

We finally consider how the membrane conformations and the energy gain Δ*E* of cooperative wrapping in tubules are affected by the length parameters of our model, i.e. by the particle radius *r*_*p*_, the standard deviation *σ* of the adhesion potential (2), and the minimum radius *r*_*n*_ of membrane necks, relative to the radius *r*_*m*_ of membrane segments adhering at preferred particle-membrane separation, which can be taken as reference length or “length unit” in our continuum model. In Ref. 7, the particle radius *r*_*p*_ was taken to be equal to *r*_*m*_, and the radius *r*_*n*_ of membrane necks was allowed to attain arbitrary small values. With these assumptions, the minimum-energy membrane shapes and the energy gain Δ*E* of cooperative wrapping only depend on the range of the adhesion potential relative to the membrane radius *r*_*m*_, i.e. on the ratio *σ/r*_*m*_ here, besides the dimensionless parameters *u* and *γ* defined in Eqs. (3) and (4). For tensionless membranes with *γ* = 0 as in Ref. 7, Figure 6a illustrates that the energy gain Δ*E* of cooperative wrapping in the particle system of Figure 1d modelled with *r*_*p*_ = 19 nm, minimum neck radius *r*_*n*_ = 2.5 nm, and membrane radius *r*_*m*_ = 23 nm clearly decreases when the standard deviation of the adhesion potential (2) is reduced from *σ* = 1 nm to 0.5 nm and 0.25 nm. In contrast to Ref. 7 the energy gain vanishes, i.e. becomes non-negative, at large values of *u* for reduced potential ranges with *σ* = 0.5 nm and *σ* = 0.25 nm. Because the particles attain distances *d* in tubules clearly larger than the minimum value 2*r*_*p*_ determined by the particle radius *r*_*p*_ at large values of *u*, this difference to Ref. 7 can only result from the finite minimum value *r*_*n*_ = 2.5 nm of the neck radius. With decreasing potential range, the membrane tubules attain necks with minimum radius 2.5 nm already at smaller values of *u* (see Figure 6b). The vanishing en-^0^ ergy gain at large *u* for reduced potential ranges can be understood by comparing the total energy densities *e* for the finite minimum value *r*_*n*_ = 2.5 nm of the neck radius (full lines in Figure 7) to total energy densities with unconstrained neck radii as in Ref. 7 (dashed lines in Figure 7). For the minimum neck radius *r*_*n*_ = 2.5 nm, the minima of the total energy densities at *σ* = 0.5 nm and 0.25 nm are shifted to smaller values of *z/r*_*m*_, compared to the profiles for unconstrained necks. An integration of the energy densities for the minimum neck radius *r*_*n*_ = 2.5 nm leads the values Δ*E* ≃ −0.65 *κ*, −0.25 *κ*, and 0.04 *κ* for *σ* = 1 nm, 0.5 nm, and 0.25 nm, respectively. The positive Δ*E* value for *σ* = 0.25 nm implies that the interplay of bending and adhesion energies in the contact regions at which the membrane detaches from the particles in necks is no longer favourable, compared to a corresponding spherical membrane segment *r*_*m*_ for an individually wrapped particle. An integration of the energy densities for unconstrained necks leads to the values Δ*E* ≃ −0.65 *κ*, −0.29 *κ*, and −0.14 *κ* for *σ* = 1 nm, 0.5 nm, and 0.25 nm, respectively. The resulting neck radii *r*_*n*_ = 1.51 nm and 0.80 nm for *σ* = 0.5 nm, and 0.25 nm, however, are unrealistically small. For *σ* = 1 nm, the membrane profiles are not affected by the minimum value 2.5 nm of the neck radius, because the resulting neck radius *r*_*n*_ = 2.95 nm is larger than this minimum value at which the neck is closed.

**Fig. 6.**
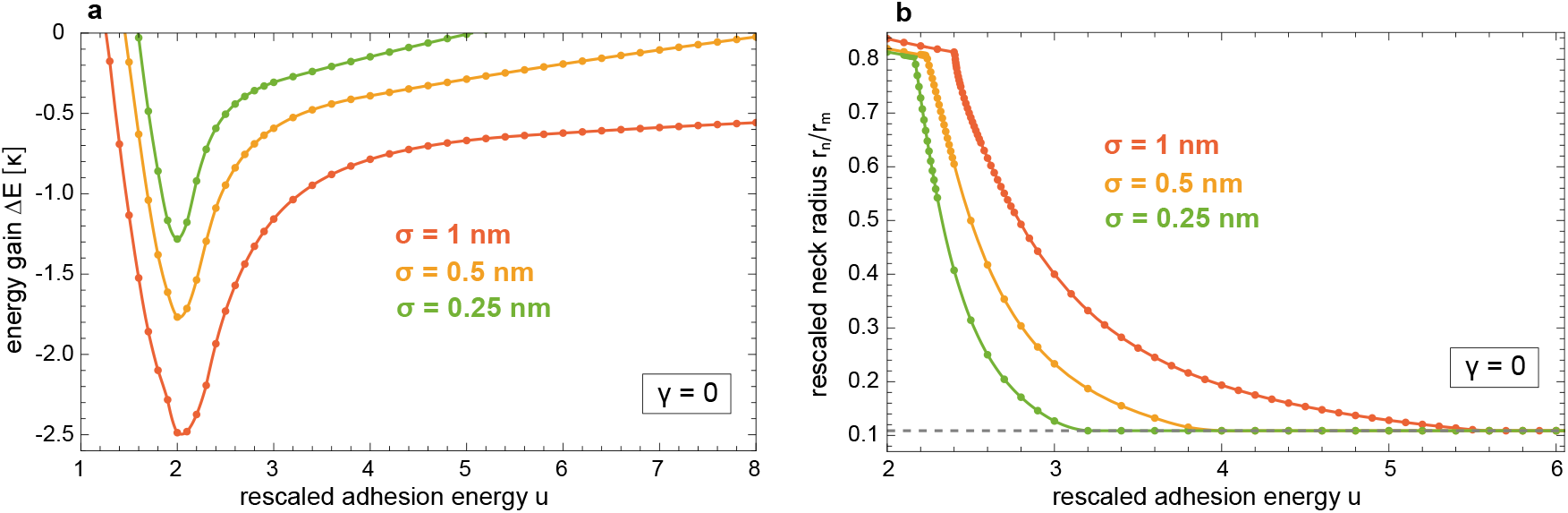
(a) Energy gain Δ*E* per central particle in long membrane tubule relative to an individually wrapped particle and (b) rescaled neck radius *r*_*n*_*/r*_*m*_ of membrane tubules versus rescaled adhesion *u* for tensionless membranes with rescaled tension *γ* = 0 and at the standard deviation *σ* = 1 nm, 0.5 nm, and 0.25 nm of the adhesion potential (2).

**Fig. 7.**
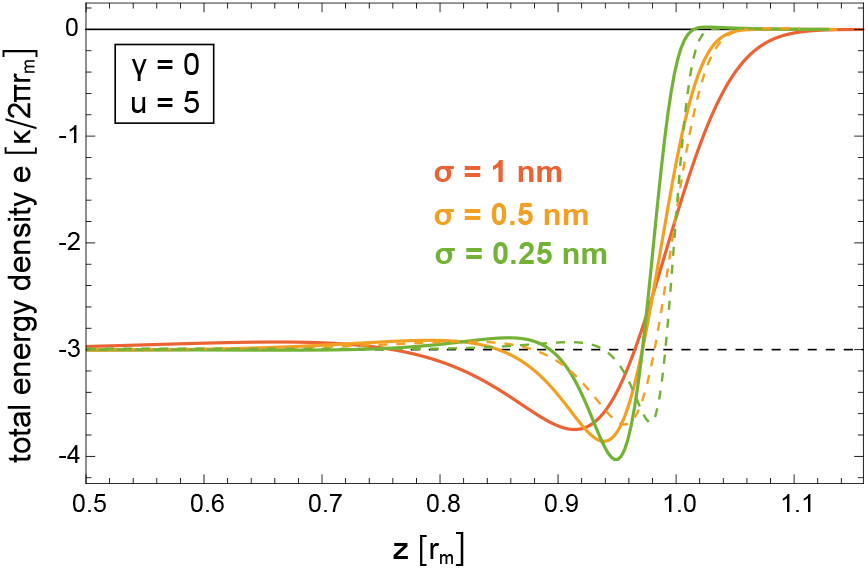
Total energy densities *e* of a central particle in a tubule in the contact region with rescaled coordinate *z/r*_*m*_ for a minimum neck radius *r*_*n*_ = 2.5 nm (full lines) and for unconstrained membrane necks (dashed lines) of tensionless membranes at the rescaled adhesion energy *u* = 5 as in Figure 6 and at different values of the standard deviation *σ* of the adhesion potential (2).

In contrast to the minimum neck radius, which affects the model results at large values of *u*, the particle radius *r*_*p*_ only affects the results at smaller values of *u* in Figure 3c at which the particles distance *d* is equal to the minimum distance 2*r*_*p*_ are only slightly larger. At larger values of *u*, the tubule conformations are not constrained by the length parameter *r*_*p*_. For an intermediate range of rescaled adhesion energies *u* in which the particle distance in the tubules is clearly larger than *r*_*p*_ and in which the radius *r*_*n*_ of membrane necks is larger than the minimum value, the model results only depend on the ratio *σ/r*_*m*_ as in Ref. 7.

## 4 Discussion and conclusions

In this article, we have extended previous modelling results ^7,19^ for the cooperative wrapping of linear chains of spherical nanoparticles in membrane tubules by considering how the minimum-energy shapes of the particle-filled tubules and the energy gain Δ*E* for the cooperative wrapping in tubules depend on the rescaled membrane tension *γ* and on various length parameters of the particles and membrane. The energy gain Δ*E* of cooperative wrapping is maximal at an intermediate value of the rescaled adhesion energy *u* at which individual particles are about half-wrapped by the membrane. For tensionless membranes, Δ*E* is maximal close to the value *u* = 2 at which the adhesion energy of a membrane segment adhering at optimal particle-membrane separation is oppositely equal to the bending energy of the segment. With increasing membrane tension, Δ*E* attains its maximal value at larger and larger values of *u* (see Figure 4b) because the membrane tension impedes wrapping, which needs to be compensated by the adhesion energy in addition to the bending energy cost of wrapping.

At large values of the rescaled adhesion energy *u*, the particles are deeply wrapped, and the energy gain Δ*E* for the cooperative wrapping in tubules is rather weakly affected by the membrane tension up to the maximal value *γ* = 1 of the rescaled tension considered here. The rather small effect of the rescaled tension *γ* on the energy gain Δ*E* at large values of *u* can be understood from the crossover length 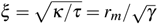 between bending-energy-dominated and tension-dominated elastic regimes. The elastic energy of the membrane is dominated by the bending energy on length scales smaller than the crossover length, and by the membrane tension *τ* on length scales larger than the crossover length ^32^. For *γ* = 1, the crossover length *ξ* is equal to the radius *r*_*m*_ of membrane segments adhering at optimal separation *lo* of the adhesion potential (2). For *γ* = 0.2 and *γ* = 0.5, the crossover length *ξ* is 5*r*_*m*_ and 2*r*_*m*_, respectively. At all these values of *γ* considered here, the crossover length *ξ* is clearly larger than the extension of the contact region in which the membrane detaches from a deeply wrapped particle. The elastic energy of this contact region, which leads to the energy gain Δ*E*, thus is dominated by the bending energy of the membrane. For the membrane radius *r*_*m*_ = 23 nm of the particle system in Fig. 1d and the typical bending rigidity value *κ* = 30*k*_*B*_*T* ^33,34^, the rescaled tension *γ* = 1 corresponds to the membrane tension 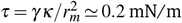, which is a relatively large membrane tension that is only about one order of magnitude smaller than the maximally sustained tension at which membranes rupture ^35^.

At small values of the rescaled adhesion *u*, the minimum total energy *E* of a particle in a membrane tubule becomes positive because the bending energy cost of the tubular membrane shape is no longer compensated by the adhesion energy gain (see Fig. 4a). Tubules, formed e.g. at smaller tension values, become instable at a rescaled tension *γ*_in_ that increases with increasing rescaled adhesion energy *u* (see Figure 8). For single particles fully wrapped by a vesicle membrane, tension-induced unwrapping has been observed after increasing the membrane tension by micropipette suction ^36^. For a partially wrapped single particle, in contrast, the minimum total energy *E* is always negative (see Fig. 4a). At small values of *u*, the bending energy of cost of partial wrapping tends to 0 with decreasing *u* because the membrane approaches a planar shape, in which a particle “sitting” on the membrane still gains adhesion energy because of the finite range of the adhesion potential (2).

**Fig. 8.**
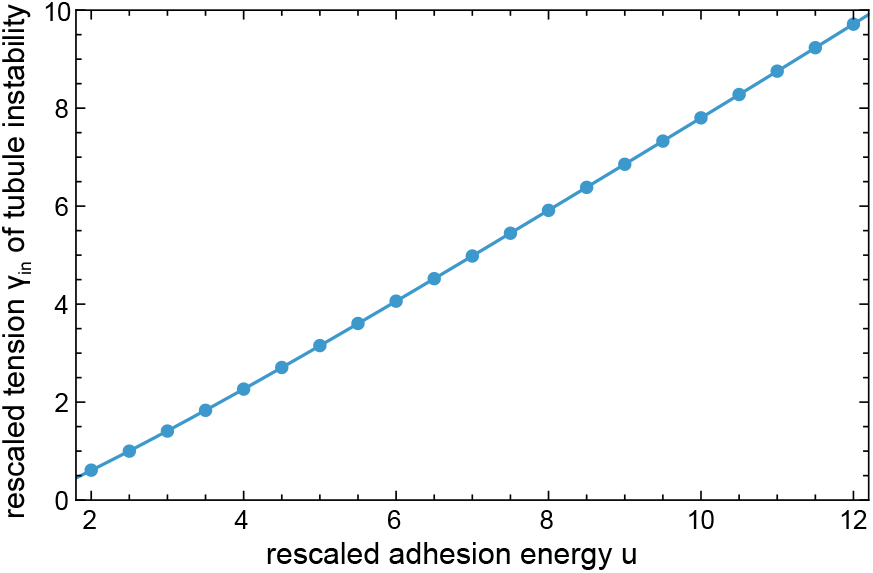
Rescaled tension *γ*_in_ at which central particles of a long tubule contribute a minimum total energy of 0 for the same parameter values as in Figure 3. At *γ*_in_, particle-filled tubules previously formed at lower rescaled tension *γ* become instable. The line represents the cubic fit function *γ*_in_(*u*) = −0.936 + 0.725 *u* + 0.0222 *u*^2^ − 0.000726 *u*^3^ of the data points.

The energy gain Δ*E* for the cooperative wrapping of spherical particles strongly depends on the range of the adhesion potential, which determines the size of the contact region in which the membrane detaches from the particle with favourable interplay of bending and adhesion energies ^7^. In addition to the potential range, another length scale that restricts the energy gain Δ*E* for cooperative wrapping at large rescaled adhesion energies *u* is the minimum radius of the membrane necks in the particle-filled tubules. For the particle system of Fig. 1d modelled with a standard deviation *σ* = 1 of the adhesion potential (2), the energy gain Δ*E* is negative and, thus, favourable also at large values of *u* for the minimum neck radius *r*_*n*_ = 2.5 nm (see Figure 6a), in line with experiments in which the adhesion energy was varied by using nanobody receptors with vastly different affinities ^19^. In these experiments, membrane tubulation was observed for all receptors with affinities beyond a threshold value.

In this article, we have focused on spherical particles adhering to initially planar membranes. For single particles, the curvature of the membrane prior to adhesion has been shown to affect the wrapping behaviour. For spherical polystyrene particles with diameters of about 1 and 2 µm that adhere to the outside of GUVs due to depletion interactions induced by polyethylene glycol (PEG) polymers in the surrounding solvent, an energetic barrier for particle wrapping has been observed ^37^, in line with theoretical calculations ^8,9^. Monte Carlo simulations of several particles adhering to vesicles show that the membrane-mediated interactions of the particles that lead to tubule formation depend on the curvature of the vesicle membrane ^38,39^ and on the area-to-volume ratio of the vesicles ^10^. Coarse-grained simulations of adhering spherical particles indicate that the membrane-mediated interactions are affected by the softness of the particles ^40^. Also for mixtures of spherical particles with different diameters and adhesion energies, cooperative wrapping has been observed in simulations and experiments ^41,42^

## Data availability

All Mathematica 14.3 notebooks used to generate and plot the data of this article are available in the open research data repository Edmond at https://doi.org/10.17617/3.OAPOLZ^43^.

## Acknowledgements

The author thanks the Max Planck Society for generous funding, and Amir Bahrami, Rumiana Dimova, Helge Ewers, Raluca Groza, Reinhard Lipowsky, and Michael Raatz for inspiring discussions in previous joint work on the article topic. The author is grateful to Erich Sackmann for his pioneering work on supported membranes ^44^ and on the specific adhesion of reconstituted vesicles by receptor-ligand complexes ^45^, which deeply affected the author’s scientific path.

